# Building a local community of practice in scientific programming for Life Scientists

**DOI:** 10.1101/265421

**Authors:** Sarah L. R. Stevens, Mateusz Kuzak, Carlos Martinez, Aurelia Moser, Petra Bleeker, Marc Galland

## Abstract

In this paper, we describe why and how to build a local community of practice in scientific programming for life scientists that use computers and programming in their research. A community of practice is a small group of scientists that meet regularly to help each other and promote good practices in scientific programming. While most life scientists are well-trained in the laboratory to conduct experiments, good practices with (big) datasets and their analysis are often missing. We propose a model on how to build such a community of practice at a local academic institution, present two real-life examples and introduce challenges and implemented solutions. We believe that the current data deluge that life scientists face can benefit from the implementation of these small communities. Good practices spread among experimental scientists will foster open, transparent and sound scientific results beneficial to society.

## Introduction

### Life Sciences is becoming a data-driven field

In the last ten years, since the advent of the first next-generation sequencing (NGS) technologies, DNA and RNA sequencing costs have plunged to levels that make genome sequencing an affordable reality for every life scientist. Yet the vast majority of wet lab biologists need tailor-made, practical training to learn scientific programming and data analysis [6–9, 13, 14]. Current efforts in bioinformatics and data science training for life scientists have been initiated worldwide to cope with these training demands [10–14].

### Good practices in scientific programming are needed to increase research reproducibility

Modern biology is facing reproducibility issues [15]. While evidence suggests this might not be as bad as it sounds [16], there is clearly a need for increased reproducibility. For instance, out of 400 algorithms presented at two conferences, only 6% had published their corresponding code [17]. Thus, most research code remains a “black box” [18] although programming is a central tool in research [19]. Use of laboratory notebooks is widely taught in biology but not emphasized for coding. Both code documentation and better practices in data management are needed so anyone can redo or understand the analyses later on. Part of the solution lies in dedicated training to researchers to promote good programming practices [20]. One of the recent relevant initiatives is the FAIR (Findable, Accessible, Interoperable and Reusable) principles initiative which provides guidelines to boost reproducibility and reuse of datasets [21]. Therefore, the long term goal of any programming scientist should be to steward good practices in code-intensive research by promoting open science, reproducible research and sustainable software development.

### Part of the solution: building a local community of practice

Training workshops in scientific programming are often offered as one-time courses but researchers would benefit from a more permanent support. Fueled by Etienne Wenger’s idea that learning is usually a social activity [22, 23], we propose to build a local community of practice in scientific programming for life scientists. This community fulfills the three requirements of Wenger’s definition: it has a specific domain i.e. bioinformatics and data science, its members engage in common activities e.g. training events, and they are practitioners i.e. researchers currently engaged in research that involves scientific programming. Community building and organization is a field in itself that has been considerably reviewed [24–29]. Requirements include a few motivated leaders and a safe environment where participants can experiment with their new knowledge [26]. As stated by Wenger and Snyder [30], communities of practice “help to solve problems quickly”, “transfer best practices” and “develop professional skills”. While short-term immediate issues (”help me now to debug my code”) can be solved, the community also has the capacity to steward solutions for long-term data-related problems (”how do I comply with the FAIR guidelines?”) and can therefore help to solve reproducibility issues. Communities of practice can also foster the adoption of good practices [31] since by co-working with their peers, scientists are probably more likely to compare their methods and embrace best practices.

This paper will explicitly describe why and how to build a local community of practice in scientific programming. We propose a model of how to build such a community that we exemplify in two case studies. Finally, we discuss the challenges and possible solutions that we encountered when building these communities. Overall, we believe that building these local communities of practice in scientific programming will support and speed-up scientific research, spread good practices and, ultimately, help to tackle the data deluge in the life sciences.

## Why do we need to build up local community of practice in scientific programming?

### Isolation

Wet lab biologists are increasingly being asked by their supervisors to analyze a set of pre-existing data in labs where their peers have little to no coding experience. Without access to experienced bioinformaticians, they can lead to a sentiment of isolation deleterious to their work.

### Self-learning and adoption of bad practices

In such a scenario, most researchers tend to invent their own solution sometimes reinventing the wheel. While wasting time, it also leads to the adoption of bad practices (lack of version control) and irreproducible results. While some compiled easy-to-use software such as samtools [32] can help to get started, typically researchers need to build their own collection of tools and scripts. For instance, version control is essential: we believe that using git^1^ and github^2^ for instance should be considered a mandatory, good practice just like accurate pipetting in the molecular lab.

### Apprehension

Researchers may also fear the breadth of knowledge they need before achieving anything which may lead to “impostor syndrome“: the researcher feels like he will be exposed as a fraud and someone more competent will unveil his lack of knowledge of coding and bioinformatics. This also inhibits continued learning since the researcher is then afraid to ask for help.

### The issue of how to get started

Learning to code in a research team is akin to an apprenticeship. The ‘apprentice’ will benefit from the experience and knowledge of more experienced team members. For instance, a researcher working on RNA-Seq for several years will be able to demonstrate the use of basic QC tools, short-read aligners, differential gene expression calls, etc. Yet, many research teams do not have an experienced bioinformatician on staff. Even in the best case where an expert bioinformatician is available, it may be problematic for beginners to get all their knowledge in one field from one person. Instead, we propose that building a community to spread good practices and help to connect novices and experts. Ideally, a novice should make progress toward increased skill levels, as illustrated in Fig 1 [33].

**Fig 1.**
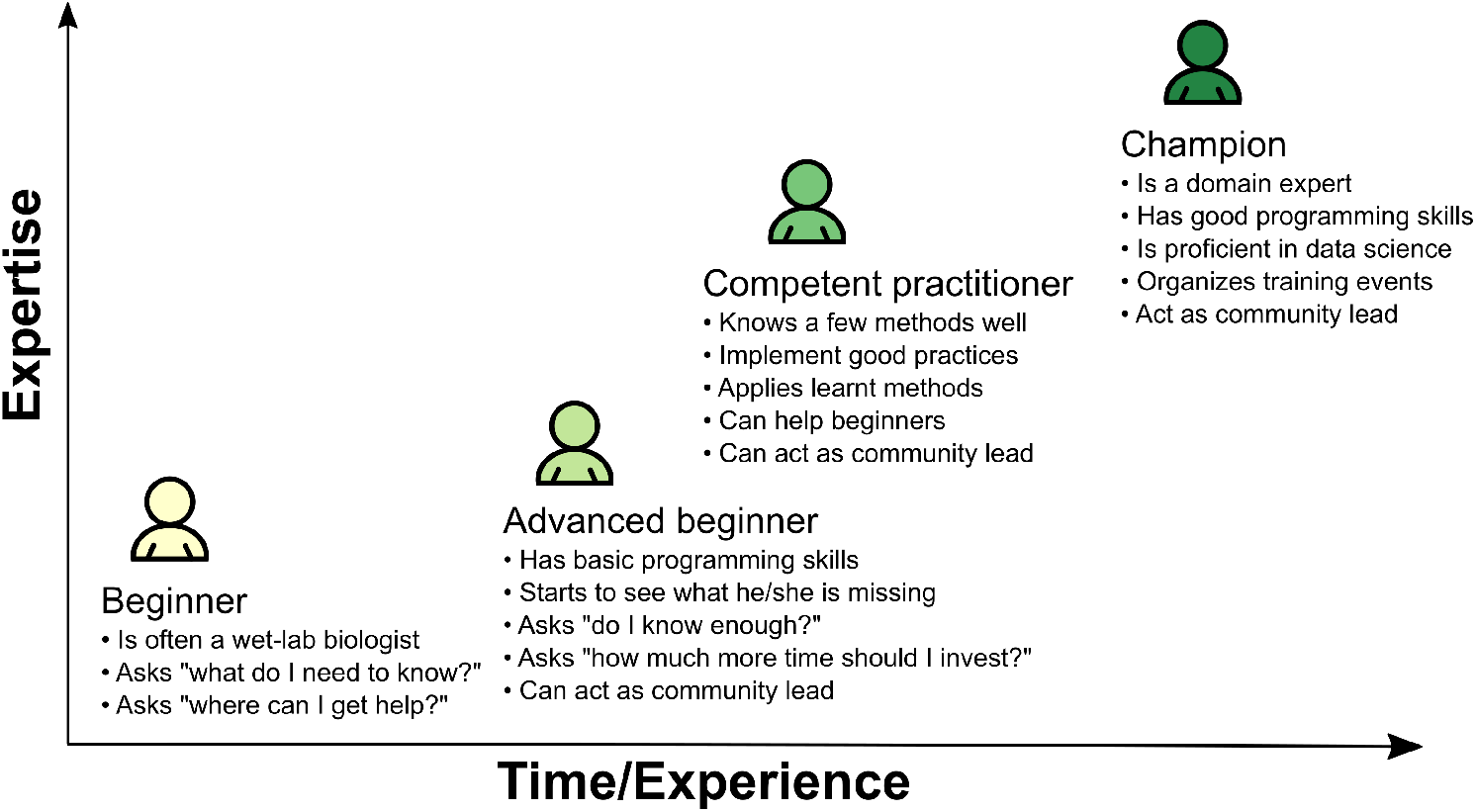
Different learning stages in scientific programming. This figure displays the different stages of learning encountered by experimental biologists.

## How do we build local communities in scientific programming? A model inspired by experience

Here, we propose a three-stage working model (Fig 2) to create a local community of practice in scientific programming composed of life scientists at any given institution without any prior community structure.

**Fig 2.**
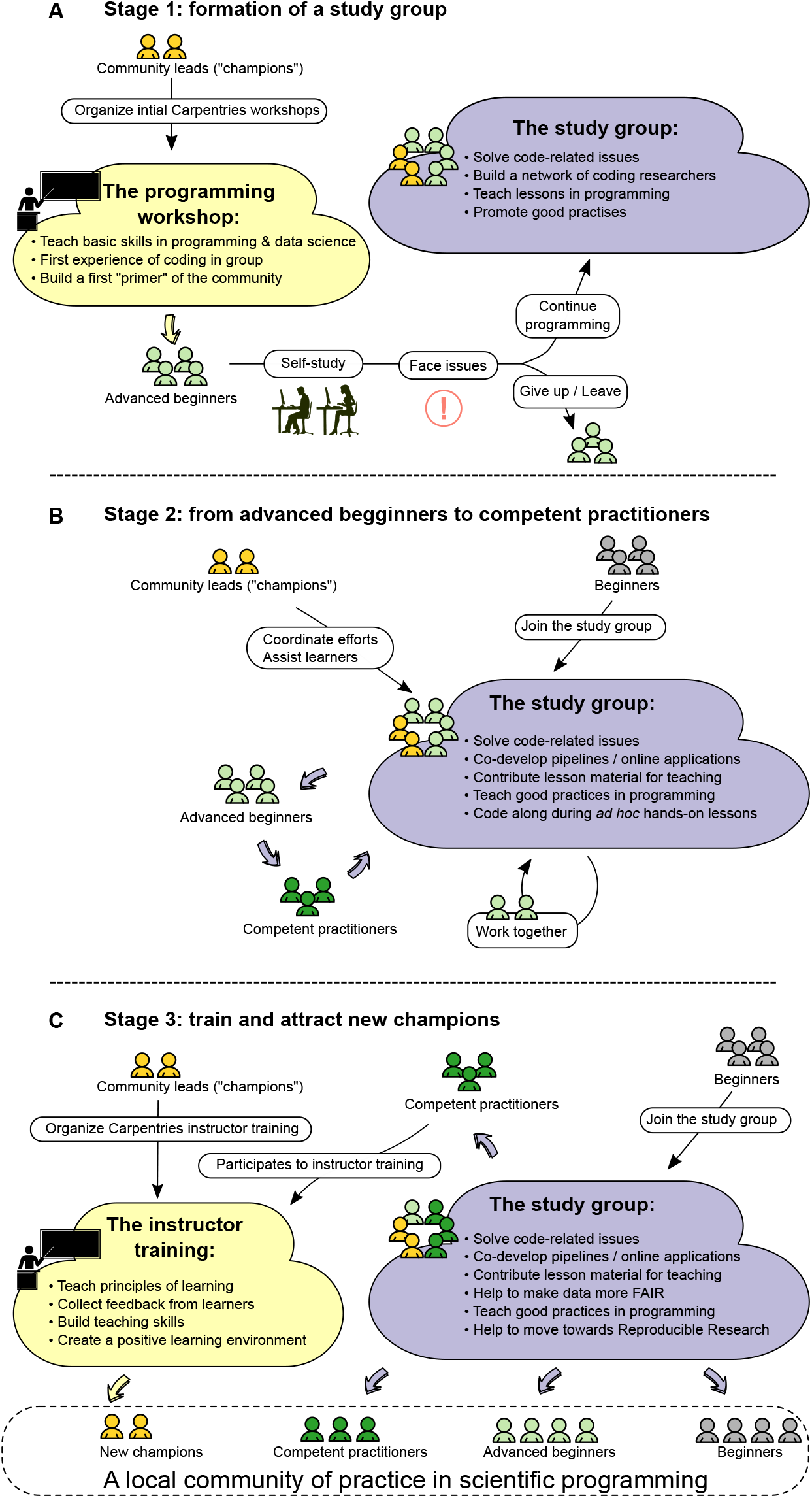
(Previous page.)A three-step model to build a local community of practice in scientific programming for life scientists. (A) First, a few scientists acting as community leads set up one or more Carpentries workshops to impart basic programming and data science skills to wet lab life scientists. After completion of the workshop, the novices will often face programming issues that need to be solved frequently. Furthermore, they need to continue to learn new programming skills. Therefore, a local study group such as a Mozilla Study Group can be formed by community leads (”champions”) and “advanced beginners” to foster a regular meeting place for solving programming issues together and discovering new tools. (B) By attending a regularly scheduled study group, advanced beginners start to work together and make progress. Together with additional guidance and *ad hoc* assistance by community leads, some advanced beginners become “competent practitioners”. (C) Finally, as some “competent practitioners” attend the Carpentries’ instructor training sessions, new community leads (”champions”) are trained. In addition, the local study group keeps attracting new beginners. Study group sessions together with optional Carpentries events help to educate community members and help them to become “advanced beginners” and “competent practitioners”. As “competent practitioners” become community “champions”, this closes the loop and help the local community of practice become fully mature with all categories of learners present.

In stage 1, we form the “primer” of a local community of practice by first running basic programming workshops organized by local community leads (”champions”) and then coupling them to formation of a study group. Champions do not necessarily have to be experts themselves. In our experience, Carpentries workshops work well since they provide training aimed at researchers and possess a long history of teaching programming to scientists [11, 20]. These programming workshops serve as a starting point for both learning and gathering researchers together in one room where people are actively paired and invited to learn about each other. Often beginners and bioinformaticians who might have never met despite working at the same institution will connect and engage afterwards.

When absolute beginners join these workshops, they become “advanced beginners” once they gain some programming notions. During their daily work, “advanced beginners” often lack the support needed to face programming issues that they may encounter frequently. Community “champions” and “advanced beginners” can “seed” a local community of practice (Fig 2) which meet regularly to continue practicing the skills they learned at these programming workshops. Therefore, a local co-working group that follow a well documented handbook such as that of the Mozilla Study Group^3^ should be set-up with a regular meeting schedule. Other forms of co-working groups can be used but we believe that Mozilla Study Groups offer the best existing model.

In stage 2, the study group becomes a regular practice for advanced beginners where they progressively become competent practitioners (Fig 2). This study group also welcomes new novice members as they join the research institution or as they hear about the existence of the group. The community leads will provide guidance, specific lessons, and assistance during hands-on practicals which will nurture the community and raise the community global scientific programming level. Again, leading sessions is not restricted to champions and any motivated individual can lead. Also, champions do not necessarily have to be experts themselves but can instead invite experts and facilitate discussions. At the end of this stage, most advanced beginners will likely have become competent practitioners.

In stage 3, a subset of the competent practitioners from the local community will become community leads (”champions”, Fig 2) by increasing their teaching and facilitating skills and recognizing the skill level of their audience (Fig 1). These competencies can be attained by becoming a Carpentries instructor which requires attending an instructor training event: these sessions can be organized by initial community champions since they usually have both the network and know-how to set-up these specific workshops. Once again, it is not mandatory to rely on the Carpentries Foundation organization as long as competent practitioners get a deeper knowledge of teaching techniques where they improve their own skills. However, we now have a good perspective on the long-term experience and success of the Carpentries Foundation with over 500 workshops organized and 16,000 attendees present [11, 12].

## Case studies

### The Amsterdam Science Park example

In October 2016, Mateusz Kuzak, Carlos Martinez and Marc Galland organized a two-day Software Carpentry workshop in Amsterdam to teach basic programming skills (Shell, version control and Python) to a group of 26 wet lab biologists. This started a dialog about the skills life scientists need in their daily work. After a few months, a subset of the workshop attendees made progress but most of them did not continue to program either because (i) they did not need it at the time, (ii) they felt isolated and could not get support from their peers or (iii) they did not make time for practice alongside regular lab work. Thus, a regular meetup group was needed so that researchers with different programming levels could help and support each other. Hence, in March 2017, we started up the Amsterdam Science Park Study Group following the Mozilla Study Group guidelines. We quickly decided to stick to the guidelines suggested by the Mozilla Science Lab^4^. Originally, we started with one scientist from the University of Amsterdam (Marc Galland) and two engineers in software engineering (Mateusz Kuzak and Carlos Martinez). But after five months, we decided to gather more scientists to build up a community with expertise in R and Python programming as well as from different scientific fields (genomics, statistics, ecology). Most study group members came from two different institutes which helped the group to be more multidisciplinary. At the same time, a proper website^5^ was set-up to streamline communication and advertise events.

### The University of Wisconsin-Madison example

At the University of Wisconsin-Madison, Sarah Stevens started a community of practice in the fall of 2014 centered around Computational Biology, Ecology and Evolution (”ComBEE”). It was started as a place to help other graduate students to learn scientific coding, such as Python and discuss scientific issues in computational biology, such as metagenomics. The main ComBEE group meets once a month to discuss computational biology in ecology and evolution. Under the ComBEE umbrella, there are also two spin-off study groups, which alternate each week so that attendees can focus on their favorite programming language. Later in ComBEE’s development, Sarah transitioned to being a part of the Mozilla Study Group community, taking advantage of the existing resources to, for instance, build their web page^6^.

Early in the development of ComBEE, the facilitating of the language-specific study groups was delegated on a semester by semester basis: this helped to keep more members involved in the growth and maturation of the local community. One of the early members of ComBEE was a life sciences graduate student who had recently attended a Software Carpentry Workshop and had no other experience doing bioinformatics. He wanted to continue his development and was working on a very computationally intensive project. He has since run the Python Study Group for several semesters and is now an exceedingly competent computational biologist. He continued to contribute back to the group through the end of his PhD, lending his expertise and experience to the latest study group discussions. The ComBEE study group is now more than three years old and acts as a stable resource center for new graduate students and employees.

## Room for improvement: challenges and solutions learned from experience

Below we describe essential components of a successful community of practice based on both literature [25–27, 29] and experience.

### Gather a core group of motivated individuals

One of the first tasks for setting up a community of practice is to gather a team of motivated individuals that will act as leaders of the community [26, 27]. To recruit these leaders, one can:

- Rely on existing communities e.g. “R lunch group” since these informal groups are often lead by motivated individuals.
- Recruit scientists that share similar values such as:

– Advocating Open Science
– Having a collaborative attitude
– Show tolerance towards cultural and scientific differences
– Being supportive of beginners and lifelong learners in general
- Search within institutions with a reasonably big size e.g. Universities.

### Keeping participants coming and engaging into the community

For someone who is part of the “core team” of a study group, the challenge is to attract experts or new members and ensure that they regularly participate in activities (lessons, co-working sessions, organizational meetings) [26, 27, 29]. Among possible incentives to keep new members and leaders engaging, we suggest to tell them that they can:

- Reach out to a wider audience by participating to lessons, workshops, etc.
- Improve their teaching skills and eventually become a Carpentries instructor
- Solve basic issues for several beginners simultaneously through workshops
- Lead the community for a semester and thereby develop their leadership
- Tailor topics to their interests
- Increase their group management, communication and networking capacities

### How to deal with the ever-ongoing turnover at academic institutions

The constant turnover of students and temporary staff remains a continual challenge. Keeping the local community ongoing requires a critical mass both for the core team and for the audience. Yet, the high turnover of students and staff also has its positive sides: a dynamic environment brings in new people eager to learn and with relevant knowledge to share in the group. We recommend using the turnover of people to your advantage by making an effort to recruit both new members and champions. Some practical solutions include:

- Advertising the community through its leaders: people bring people through word to mouth
- Invite permanent staff to sustain the community development
- Use the turnover to your advantage: quickly invite newcomers to join the community

### Dealing with the impostor syndrome

Creating a safe learning environment is one of the requirement for a thriving community of practice [26]. To encourage beginners and newcomers to participate and feel welcome, we recommend to:

- Enforce a Code of Conduct following an existing example^7^ to set-up expectations and promote a welcoming atmosphere
- Promote all questions and forbid surprise reactions to very basic questions (”What is the Shell?”, “Oh you don’t know?”)
- Ban in-depth technical discussions that alienate novices

### Community leadership and institutional support

An effort should be made to assign clear and specific roles to administration members of the local community based on their expertise and interest. Another challenge is to secure funding and people support from the local institution [26, 27]. To do so, we advise to:

- Delegate as much as possible to promote leadership: appoint someone to lead the community for a semester for instance
- Get support from the local institution as soon as possible in terms of money, time and/or staff

### Community composition

Another important aspect to consider is the composition of the community. We have identified the following types of community members as common components of the community:

- Absolute and advanced beginners: these are people with the most basic level of knowledge. For them, the motivation to be part of a community is obvious: they want to learn programming and often need rapid assistance to complete their research.
- Competent practitioners: these are people who already competent in a particular bioinformatics/data science domain. For them, contributing to the community is a good way to reinforce their set of talents. Often, competent practitioners make excellent teachers, as they are able to easily relate to the beginner state of mind. In turn, this increases their learning and teaching skills.
- Experts: these are people with the highest experience level on a particular skill. Experts usually reinforce their knowledge by ‘going back to basics’: it is useful for them to understand what are the usual *gotchas for novices*. Building a local community of practice provides experts with an opportunity to help novices in a more structural way instead of helping each one individually.

### Practical considerations

In our experience, we have found the following practical tips to be useful:

- Gather a critical mass of at least 10 recurrent community members that regularly attend meetings and community sessions
- Send meeting notifications in advance and frequently enough: schedule the meetings well-in-advance and keep a consistent day, time and place to help people remember them.
- Have weekly or fortnightly meetings so that it is a compromise between researchers’ schedules and community development.
- Organize meetings in a relatively quiet environment with a good Internet connection. Places such as a campus café outside of busy hours or a small conference room can be good places to start and help to keep an informal and welcoming atmosphere,

## Conclusion

We hope that our model and the lessons learned from our experience described in this paper will save time and effort for future community leads when they start their own local community of practice in scientific programming. Building such a community is far from trivial and we, as scientists, are perhaps not the most proficient on community building and organization [24–28]. Since “progress will not happen by itself” [20], a community of practice in scientific programming will bring many benefits to its members and to their institution: it fosters the development of new skills for its members, breaks down “mental borders” between departments, networks domain experts at a local site and helps to retain knowledge that would otherwise be lost with the departure of temporary staff and students.

The convergence of the “big data” avalanche in biology and new FAIR requirements for data management [21] makes it more and more important for wet lab researchers to conduct good scientific programming, efficient data analysis, and proper research data management. Eventually, these local communities of practice in scientific programming should speed up code-intensive analyses, promote open science, research reproducibility and spread good practices at a given institution.

## Acknowledgments

We are thankful to the Carpentries Foundation for assistance in workshop organization. We kindly acknowledge the Mozilla Foundation for assistance in starting and maintaining the Study Groups. We would also like to thank the members and leaders of the Amsterdam Science Park Study Group and that of the Computational Biology, Ecology and Evolution (ComBEE) Study Group and the Carpentry community at the University of Wisconsin-Madison.

## Supporting Information

<u>Galland+blurb+RR.docx</u>

## Footnotes

1 https://git-scm.com/

2 https://github.com/

3 http://mozillascience.github.io/studyGroupHandbook/

4 https://mozillascience.github.io/study-group-orientation/

5 https://scienceparkstudygroup.github.io/studyGroup/

6 https://combee-uw-madison.github.io

7 https://docs.carpentries.org/topic_folders/policies/code-of-conduct.html

